# Higher-order dialectic variation and syntactic convergence in the complex warble song of budgerigars

**DOI:** 10.1101/2022.09.18.508412

**Authors:** Abhinava Jagan Madabhushi, Nakul Wewhare, Priya Binwal, Anand Krishnan

## Abstract

Dialectic signatures in animal acoustic signals are key in identification and association with group members. Complex vocal sequences may also convey information about behavioral state, and may thus vary according to social environment. Many bird species learn and modify their complex acoustic signals throughout their lives. However, the structure and function of vocal sequences in perennial vocal learners remains understudied. Here, we examined vocal sequence variation in the warble song of budgerigars, and how these change upon contact between social groups. Budgerigars are perennial vocal learners which exhibit fission-fusion flock dynamics in the wild. We found that two captive colonies of budgerigars exhibited colony-specific differences in the syntactic structure of their vocal sequences. Individuals from the two colonies differed in the propensity to repeat certain note types, forming repetitive motifs which served as higher-order signatures of colony identity. When the two groups were brought into contact, their vocal sequences converged within months of contact, and these colony-specific repetitive patterns disappeared, with males from both erstwhile colonies now producing similar sequences with similar syntactic structure. Our data suggests that budgerigars can encode substantial information in the higher-order temporal arrangement of notes/vocal units, which is modified throughout life by social learning as groups of birds continually associate and dissociate. Our study sheds light on the importance of examining signal structure at multiple levels of organisation, and the potential for psittaciform birds as model systems to examine the influence of learning and social environment on acoustic signals.

## Introduction

Animal sensory signals are often complex and elaborate, conveying a variety of information such as behavioral context, mate quality, presence of a predator, territoriality and group membership (Bradbury and Vehrencamp, 2011). Diverse animal taxa employ sound in the form of acoustic signals to communicate both short- and long-range information. Complex signals support communication of contextual information, by changes in the content and structure of signals (Bradbury and Vehrencamp, 2011; Engesser et al., 2019; Hebets and Papaj, 2005; Suzuki et al., 2018). For example, members of a species may change the structure of a single element in a signal, or the temporal structure (syntax) of an acoustic signal sequence to convey information (Bhat et al., 2022; Bohn et al., 2009; Engesser et al., 2019; Leroux et al., 2021; Suzuki et al., 2018; Zuberbühler, 2018). The larger the repertoire of elements, the greater the combinatorial diversity of sequences, and the more complex the information content of signals (Backhouse et al., 2022; Balsby et al., 2017; Clay and Zuberbühler, 2011; Dalziell and Welbergen, 2016; Leroux et al., 2021; Mitoyen et al., 2019; Scholes III, 2008). In addition to behavioral state, group-living animals may use complex signals to communicate group or individual identity (Mammen and Nowicki, 1981; McComb et al., 2000; McCowan and Hooper, 2002). The dialects of passerine songbirds represent one such form of variation, with individuals responding more strongly to their local dialect than to an unfamiliar one (Briefer et al., 2013; Dahlin and Wright, 2009; Marler and Slabbekoorn, 2004; Marler and Tamura, 1962; Searcy et al., 1981). Oscine passerines, in addition to certain other bird groups such as parrots, learn their complex vocal signals. Therefore, both the individual vocal units (notes or syllables) and the syntactic structure of the song are culturally transmitted across populations, resulting in dialectic variation among different populations. In passerines, primates and cetaceans, the structure of song carries signatures of population, species and individual identity (Allen et al., 2018; Deecke et al., 2010; Marler and Tamura, 1962; Mitani and Marler, 1989; Peters et al., 1980; Rendell and Whitehead, 2003).

Unlike most well-studied passerine systems, members of the parrot order Psittaciformes are perennial vocal learners, modifying their vocal repertoires throughout their lives. Various parrot species are highly social, and can rapidly modify their vocalizations, which is proposed to mediate fission-fusion dynamics in flocking species (Berg et al., 2012; Dahlin et al., 2014; Hile et al., 2000; Hile et al., 2005). For example, when smaller flocks aggregate into larger ones, or individuals move between flocks, the acoustic structure of individual flight call notes changes to match that of the new group (Dahlin et al., 2014; Farabaugh et al., 1994; Hile and Striedter, 2000; Hile et al., 2000; Hile et al., 2005; Wanker et al., 2005). This modification or vocal convergence is hypothesized to mediate affiliative and aggressive interactions between different flock members, or in pair-bonding and pair-bond maintenance (Dahlin et al., 2014; Hile et al., 2000; Hile et al., 2005; Mammen and Nowicki, 1981; Tyack, 2008). Similar convergence is observed in primates, and is thought to serve similar social functions (Candiotti et al., 2012; Elowson and Snowdon, 1994; Mitani and Gros-Luis, 1998). However, the individual call notes of parrots are often emitted as part of highly complex vocal sequences (Balsby et al., 2017; Farabaugh et al., 1992; Zdenek et al., 2015). Although the studies discussed above have identified convergence in the acoustic structure of individual call notes due to pair-bonding or mixing of flocks, the vocal sequences of parrots remain sparsely studied. In particular, there is no information on how the syntactic structure of vocal sequences are modified by social context, and whether group identity is conveyed by higher-order information within sequences as compared to individual notes. Further, there is almost no information on how the structure of vocal sequences changes when different groups come into contact.

The budgerigar (*Melopsittacus undulatus*) is a social psittaciform species native to Australia, which has been extensively domesticated for over a century. Male budgerigars emit a complex, lengthy sequence of vocalizations, called a warble song (Fig 1A), which is sung both to other males as well as females (Brockway, 1964; Farabaugh et al., 1992). Although individuals from different colonies differ in the acoustic structure of individual call notes, which converge when introduced into a new social group (Dahlin et al., 2014; Farabaugh et al., 1994; Hile and Striedter, 2000), the sequence of the warble song has received very little study in these contexts. Here, we examine the syntactic structure of the warble song in laboratory colonies of budgerigars to examine whether the complexity and syntax of the warble varies according to social context and group identity. Specifically, we test whether warble syntax differs between male-male or male-female contexts, and also whether syntactic structure encodes the identity of the colony from which the male originates. Next, we examine whether the warble song exhibits syntactic convergence when groups of birds are brought into contact. Our study thus examines the presence of higher-order dialectic variation and vocal convergence in psittaciform birds, which has implications for our understanding of complex communication in this sophisticated signaling system. In addition, perennial vocal learners such as budgerigars provide valuable insight into how other complex acoustic signals (including human language) exhibit higher-order change due to social learning.

**Fig 1.**
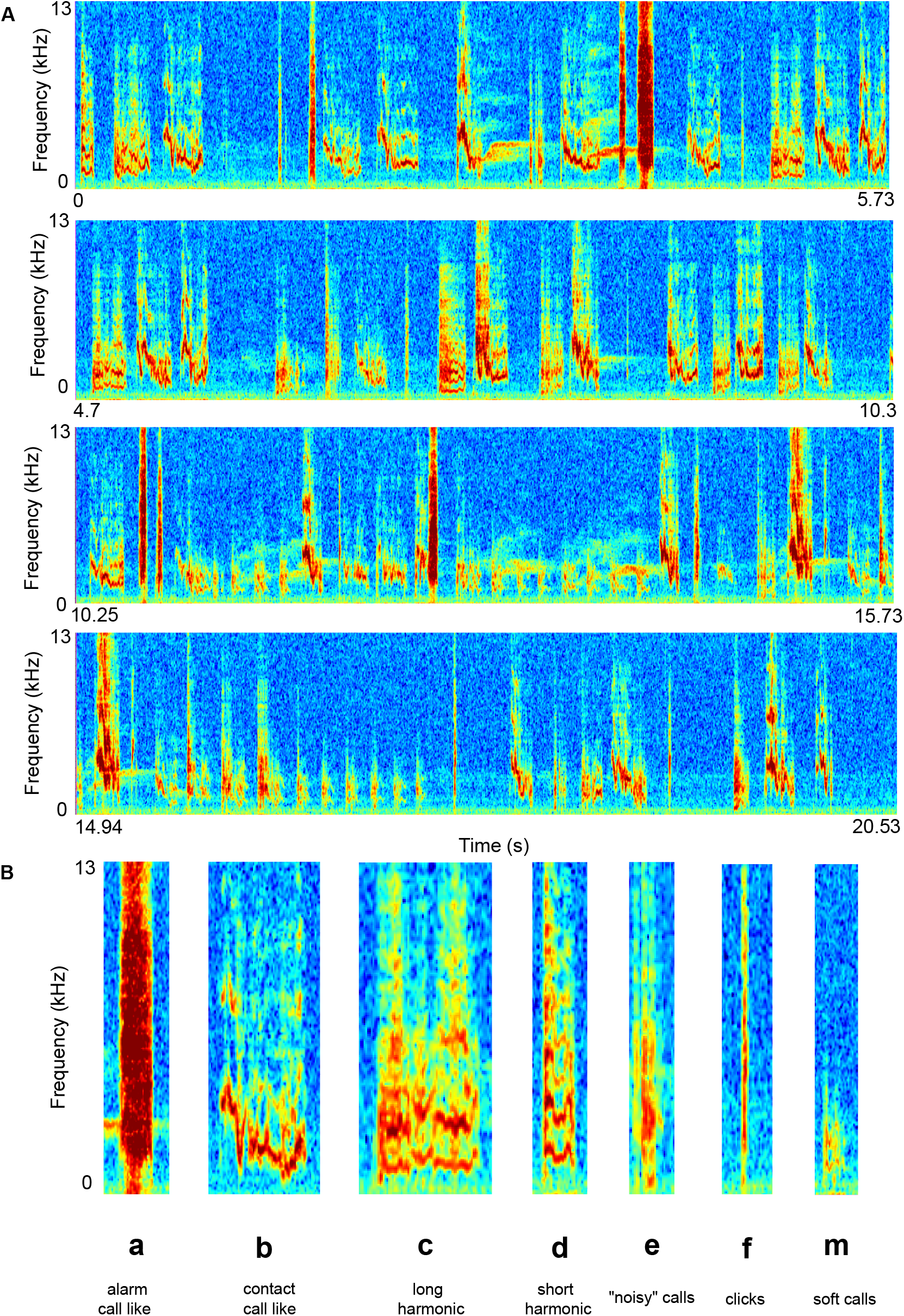
**A.** Spectrograms representing a total of 20.53 seconds of a warble song produced by a male budgerigar. **B.** Examples of the seven different note types in budgerigar warble song. a: alarm call-like elements; b: contact call-like elements; c: long harmonic sounds with duration more than 100 ms, d: short harmonic sounds with duration less than 100 ms; e: “noisy” calls - broadband sounds; f: short duration broadband ‘clicks’; m: “soft calls”-low amplitude sounds. See the Methods for further details.

## Materials and Methods

### Study Animals

We carried out our experiments at the Indian Institute of Science Education and Research (IISER) campuses in Pune and Bhopal. All procedures were approved by the Institutional Animal Ethics Committees of both institutes (Protocol Nos: IISER_Pune/IAEC/2020_01_01 and IISERB/2022/001) in accordance with the guidelines laid out by the Committee for the Purpose of Control and Supervision of Experiments on Animals (CPCSEA, New Delhi). All our experiments were carried out on captive budgerigars (*Melopsittacus undulatus*) procured from local pet stores, which were sexed by differences in cere color. In our study, Colony 1 consisted of two males and four females procured in March 2020, and Colony 2 consisted of three males and two females procured from a different dealer in August 2021. Thus, the two colonies originated from different sources. We housed the birds in separate rooms within the same lab at IISER Pune for the first part of the study, thus ensuring no visual or acoustic contact between them. We provided the birds with ad libitum water and commercially available bird seed during all experiments, and housed them under a 12-hour: 12-hour light: dark cycle. In late December 2021, the bird colony moved to IISER Bhopal, and the two colonies were combined into the same cages and housed together during and after this move. This enabled us to mimic the fission-fusion dynamics of natural budgerigar flocks and examine its effect on vocal sequences and syntax.

### Experimental Procedure and Recording

Colony 1 contained one possible male-male pair and eight possible male-female pairs (based on the number of possible combinations of individuals within a colony), whereas Colony 2 possessed three possible male-male and six possible male-female pairs. In order to examine whether social context shaped the sequences of notes emitted during warble song, we placed each of these within-colony pairs inside a soundproof acoustic enclosure of dimensions 75cm x 75cm x 75cm (Newtech Engineering Systems, Bengaluru, Karnataka, India). This enclosure was acoustically isolated from the external environment. Birds remained in the experimental enclosure for 24 hours, to habituate them to the experimental paradigm. We then recorded budgerigar vocalizations for the next 24 hrs at a sampling rate of 44.1 kHz, using Audiomoth recorders (Hill et al., 2019). The recordings were carried out in September and October 2021. As described above, we recorded warble songs from a total of nine pairs from Colony 1 (one Male-Male and eight Male-Female pairs) and nine pairs from Colony 2 (three Male-Male and six Male-Female pairs). For the second part of the study, our goal was to examine whether syntax had changed after the two colonies came into contact with each other. Thus, we re-recorded all Male-Female pairs in June 2022, nearly six months after the colonies were brought into contact. In the intervening duration, one female from Colony 1 died of natural causes. As a result, we had two fewer male-female pairs in Colony 1 for the dataset collected after contact. We therefore analyzed additional warbles of males from the erstwhile Colony 1, to ensure that both datasets had an equal number of warbles per male.

### Analysis

We defined a warble as a vocal bout with a duration of at least one second, consisting of three or more warble elements (or notes, see below for more details on the note types). Notes separated by less than 1 second were considered to belong to a single warble bout, following definitions in the literature (Farabaugh et al., 1992; Tobin et al., 2019; Tu et al., 2011). When notes were very close together in time, we applied a separation of at least 10 ms and/or a dramatic change in note structure (see below for categories used to describe note structures) to identify the boundary between two notes. For each of the pair combinations described above, we analyzed 10 warble sequences (with a good signal: noise ratio and no background sounds), using the time stamps to ensure that these warbles occurred between 8AM and 6PM when the colonies were most active. One of the 18 pairs in the 2021 dataset emitted only a single warble in the 24-hour duration, and was thus omitted from further analyses. Our final dataset consisted of a total of 310 warble sequences across the 2021 and 2022 recording experiments.

Next, we classified warble elements into different note types, broadly following the classification key employed by multiple published studies (Farabaugh et al., 1992; Tobin et al., 2019; Tu et al., 2011). Our dataset contained six such previously described categories, listed here according to the abbreviations used in the figures - a: alarm call like elements; b: contact call-like elements (frequency modulated sounds); c: long harmonic calls (harmonic sounds with duration greater than 100 ms); d: short harmonic calls (harmonic sounds with duration less than 100 ms); e: “noisy” calls that are broadband and non-harmonic; f: clicks (extremely short broadband calls). In addition, our data contained an additional note type that we named m. These notes were generally low in amplitude, and did not fit into any of the above categories. We defined these as “soft calls” (Fig 1B, Fig S1). Because multiple authors annotated note types, we performed cross-verification (where each author annotated the same warbles independently of the others) to quantify inter-observer agreement in note classification. Each person annotated five warbles from the first dataset (2021) and five from the second dataset (2022), and we then calculated the Levenshtein distances between each pair of warbles for each author (therefore, four values for each warble). Next, we normalized the Levenshtein distances to the length of each warble sequence to obtain the error as a percentage of sequence length. The average agreement in note type classification between observers was 80.16 %, which we deemed an acceptable level of agreement to proceed with analysis of syntax.

Following the classification of notes, we quantified a number of metrics of warble structure and complexity, in order to compare them across contexts and colonies. First, we calculated the Shannon Entropy and Duration (in seconds) for each warble for each pair. The Shannon Entropy (H) of a stochastic process with n states is defined as 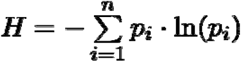. Here, *p_i_* is defined as the probability of occurrence of the *i^th^* state and 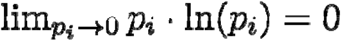. The values of H can range from 0 to ln(*n*), with greater values for H indicating that the process is more unpredictable. In the case of budgerigar vocalizations, higher values of H indicate greater variation in the note composition of warble sequences, and thus greater complexity. We then performed a Wilcoxon Rank-Sum test to compare the Shannon Entropy and Duration values across contexts and across colonies (the latter for the 2021 data, and then again for the 2022 data). Because there was only one male-male pair in colony 1, we only performed a statistical comparison of contexts for the data from colony 2. We compared social contexts for the 2021 data, and only recorded male-female pairs again in 2022 to compare the two colonies to each other before and after they were brought into contact.

Finally, we analyzed the syntactic structure of warbles, to compare how the compositions of these songs varied across colonies before and after contact. Using the note classification detailed above, we first constructed note sequences for each warble. In addition to the above described note types, and in order to incorporate the effects of silence into our analyses following a previous study (Bhat et al., 2022), we defined silent periods in the warble as instances where the inter-note interval exceeded 500 ms. This value of 500 ms was half of the duration between separate warble bouts (one second, as defined by us earlier) and also represented a period greater than the duration of all observed note types as described above. We labeled the silent periods in the warbles as ‘0’ in our note sequences.

To quantify whether certain notes tended to occur together in a warble sequence, we employed a co-occurrence metric that has been used for similar analyses in anuran vocal sequences (Bhat et al., 2022). We used this metric as it is free of assumptions about the underlying process generating vocal sequences in animal signals (for example, whether or not they follow Markov chain dynamics) (Kershenbaum et al., 2014). To elaborate, we defined ^d^C_ij_, as the probability that note type j occurs within a distance of d-1 notes of note type i. We computed ^d^C_ij_ for warbles from each pair in our dataset. To compare these observed probabilities to those expected by chance, we next constructed artificial sequences, where warble elements were randomly distributed following a stationary distribution. Here, the probability of any note type occurring at any given position in the warble was given by the proportion of that note type in the warble sequences of that colony. We computed 50 such artificial sequences to have a robust estimate of ^d^E_ij_, where ^d^E_ij_ is the probability that note type j occurs within a distance of d-1 notes of note type i due to random chance. We then computed ^d^R_ij_, defined as the ratio of ^d^C_ij_ and ^d^E_ij_ for warbles from individuals of each colony. ^d^R_ij_ is, therefore, a measure of whether note type j occurs within a distance of d-1 notes from note i more or less often than expected by random chance. A value of ^d^R_ij_ > 1 suggests that the note type j occurs within d-1 notes of note type i more often than expected by a random chance, and ^d^R_ij_ value of < 1 suggests that the note type j occurs within d-1 notes of note type i less often than expected by random chance. For the analysis described below, we used a d-value of 6, but repeated our analysis for d-values of 3 and 9 to ensure our choice of d value did not influence our findings (Bhat et al., 2022).

For the analysis above, we computed the proportion of different note types in the warbles of individuals from the two colonies. Differences in the note proportions across colonies could occur for two reasons – a) The note type occurred singly, but was emitted more or less often and b) A particular note type showed an increased or decreased tendency to repeat itself. These two possibilities result in very different outcomes in terms of the syntactic structure of sequences. To examine and distinguish between these possibilities, we computed the frequency of repeats (i.e. how often each note type occurred singly, as a repeat of two notes, and so on) for each note type in the warbles of each individual from both colonies. This enabled us to examine differences in the syntactic structure of warbles across colonies.

## Results

### Warble duration and complexity do not differ across social contexts

On average, budgerigar warbles consisted of 82.27 notes (Range: 8 - 954, total of 25,504 notes in our dataset across both 2021 and 2022 recording sessions). Warble duration did not differ across social contexts (male-male versus male-female) for the pairs in Colony 2 (Wilcoxon Rank Sum test: W=1101, n1=30, n2= 60, p-value= 0.0861) (Fig S2). Additionally, we found no difference in the Shannon Entropy values for warbles across social contexts for individuals in colony 2 (Wilcoxon Rank Sum Test: W= 812, n1=30, n2=60, p-value= 0.454) (Fig S2). As mentioned earlier, we were unable to perform a statistical comparison for colony 1 as there was only a single male-male pair in this colony. Therefore, we proceeded to examine whether warble structure (the information contained within a sequence) depended on the colony the males originated from, as opposed to the context in which they were produced. For all subsequent comparisons, we relied on male-female data to ensure the identity of the male bird producing the warble.

### Warble complexity differs across colonies and converges upon contact

Before contact, differences in warble duration between the two colonies was at the margin for statistical significance (Wilcoxon Rank Sum Test: W = 2495, p-value= 0.0654, n1=70, n2=60). Warbles produced by individuals in Colony 1 were also more complex than those produced by Colony 2 individuals, as measured by generally higher Shannon Entropy values (Wilcoxon Rank Sum Test: W= 2560, n1= 70, n2= 60, p-value= 0.0319) (Fig 2A, B).

**Fig 2.**
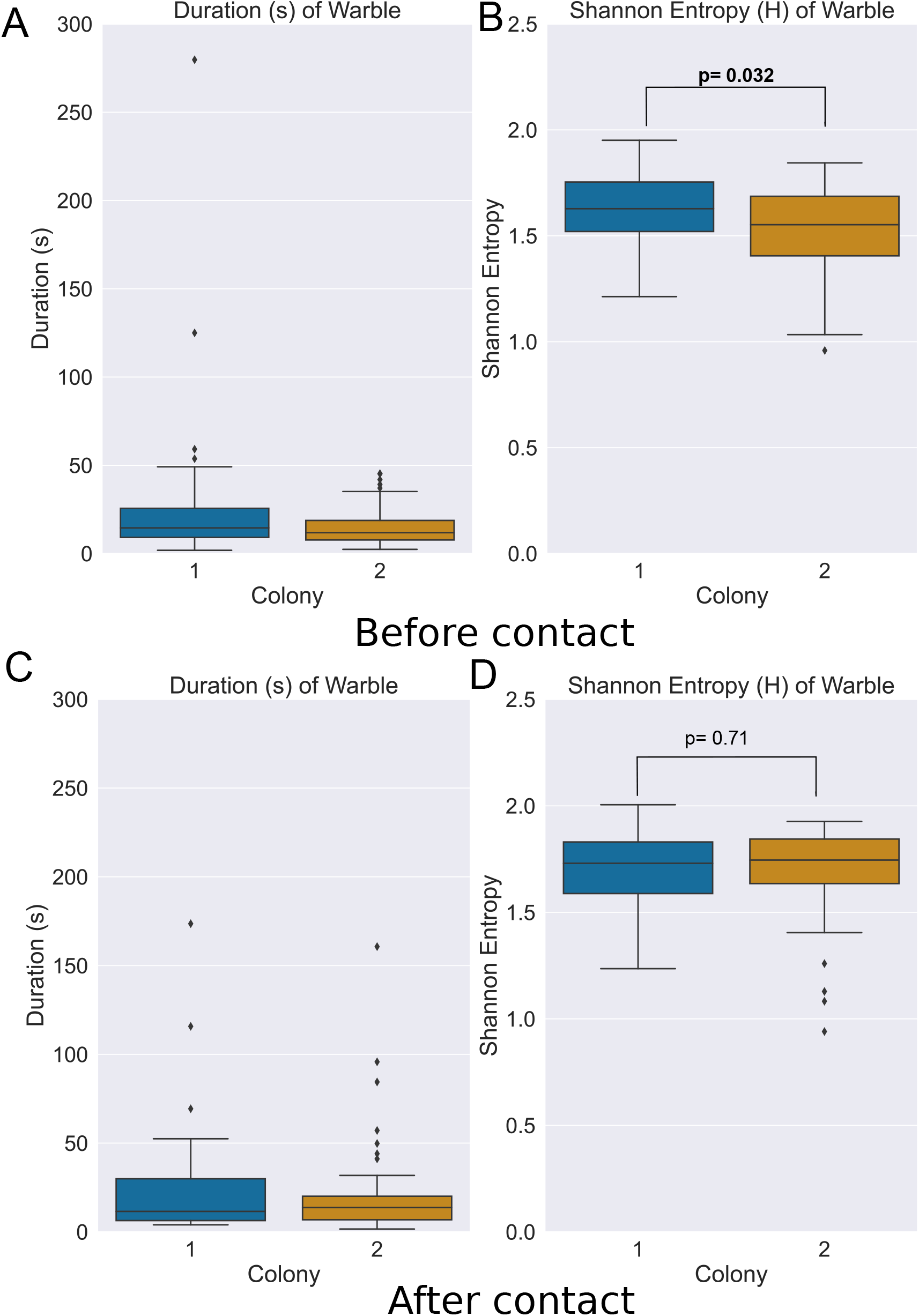
Box plots depicting the duration and Shannon entropy of warbles across the two colonies before (**A**) and after (**B**) contact. Shannon entropy for the warbles of Colony 1 differed significantly from those of Colony 2. Each data point represents one warble song, and the upper and lower edges of the box indicate the third and first quartile, respectively. The line inside each box indicates the median, and the whiskers represent 1.5 times the interquartile range. Finally, the diamonds above and below the boxes represent outliers.

After the colonies were brought into contact and recorded again a few months later in 2022, however, the differences in both duration and Shannon entropy disappeared completely (duration: Wilcoxon Rank Sum Test: W = 2402, p-value= 0.995, n1=80, n2=60; Shannon entropy: Wilcoxon Rank Sum Test: W = 2312, p-value= 0.713, n1=80, n2=60) (Fig 2C, D). Thus, the differences in warble complexity were lost when the two flocks associated with each other, mimicking natural fission-fusion dynamics. Because Shannon entropy is calculated using the proportions of different note types, our results suggested that the two colonies differed in their use of different notes within the repertoire, and that these differences were lost upon contact between colonies. When we examined the relative proportions of different note types in the repertoire of each colony before contact, we observed differences between them (Fig 3A), further supporting our assertion. Individuals in Colony 2 produced a higher proportion of d notes in particular (see Fig 1 for key). However, after contact, males from both colonies produced similar proportions of d notes, as well as all other note types (Fig 3B).

**Fig 3.**
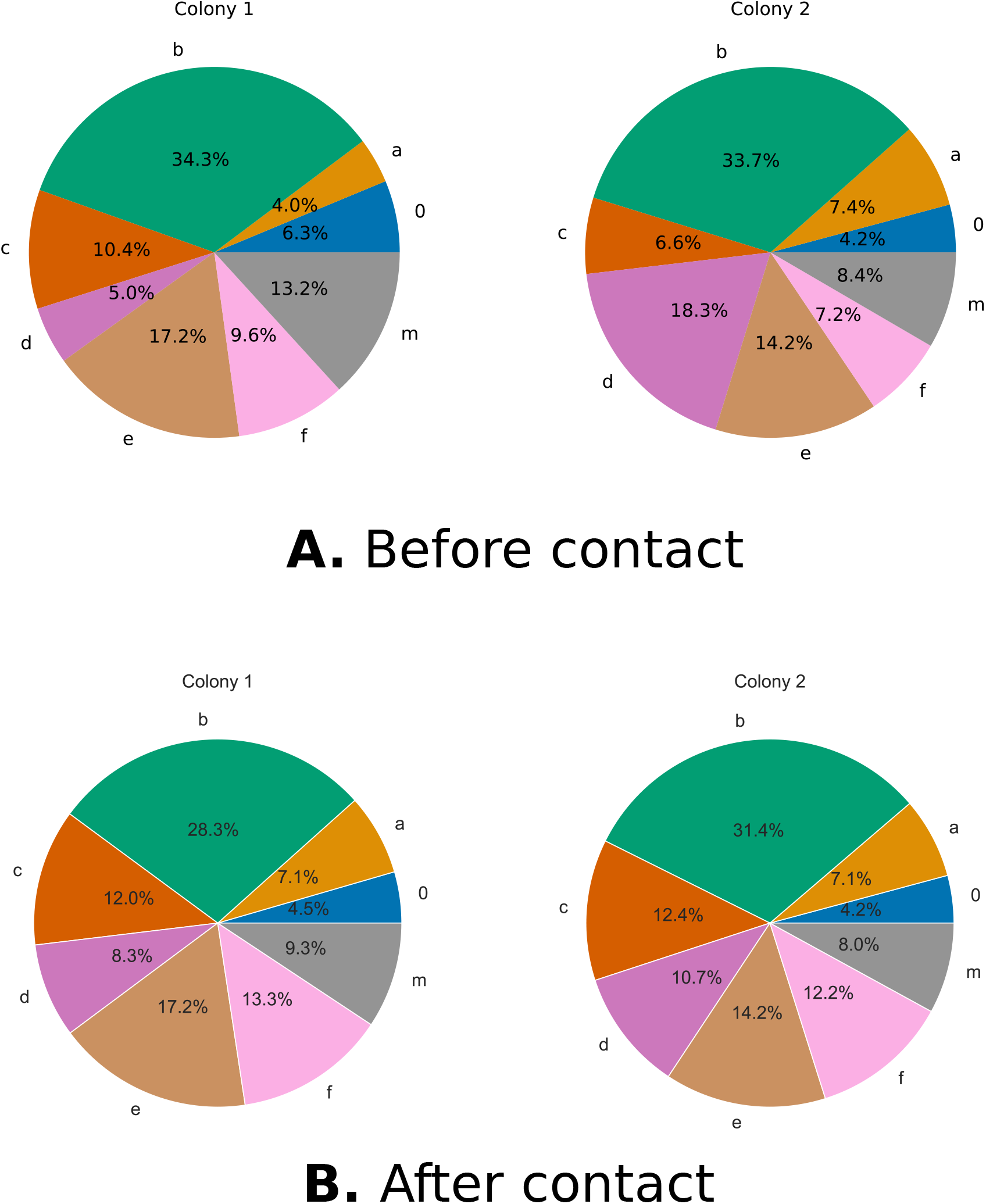
Pie chart depicting the proportions of different note types in the warbles of the two colonies before (**A**) and after (**B**) contact, as a proportion of the total notes produced. Note type d occurred in greater proportions in Colony 2 before contact, but this difference was lost within months after the colonies were brought into contact.

### Colony signatures and syntactic convergence in budgerigar warbles

To examine whether differences in warble complexity between colonies, and the subsequent changes after contact resulted from convergence of syntactic structure, we constructed note sequences for each warble. We then computed a co-occurrence matrix for warbles of each colony before and after contact. The ratio of observed to expected co-occurrence values (^d^R_ij_) was > 1 along the diagonal for both the colonies (Fig 4A). This suggests that all the note types had a propensity to repeat themselves within a warble sequence. Additionally, we also found that the off-diagonal (between notes) ^d^R_ij_ were <1. However, the tendency of d notes to repeat was higher in Colony 2 than in Colony 1. This, coupled with their higher proportion of occurrences in Colony 2 repertoire, suggested a higher-order signature of colony identity based on a tendency to repeat certain note types more often. The increased tendency for d notes to co-occur was lost after contact between colonies (Fig 4B). This pattern was independent of the distance (d-value, see Methods) used in this analysis (Fig S3, S4).

**Fig 4.**
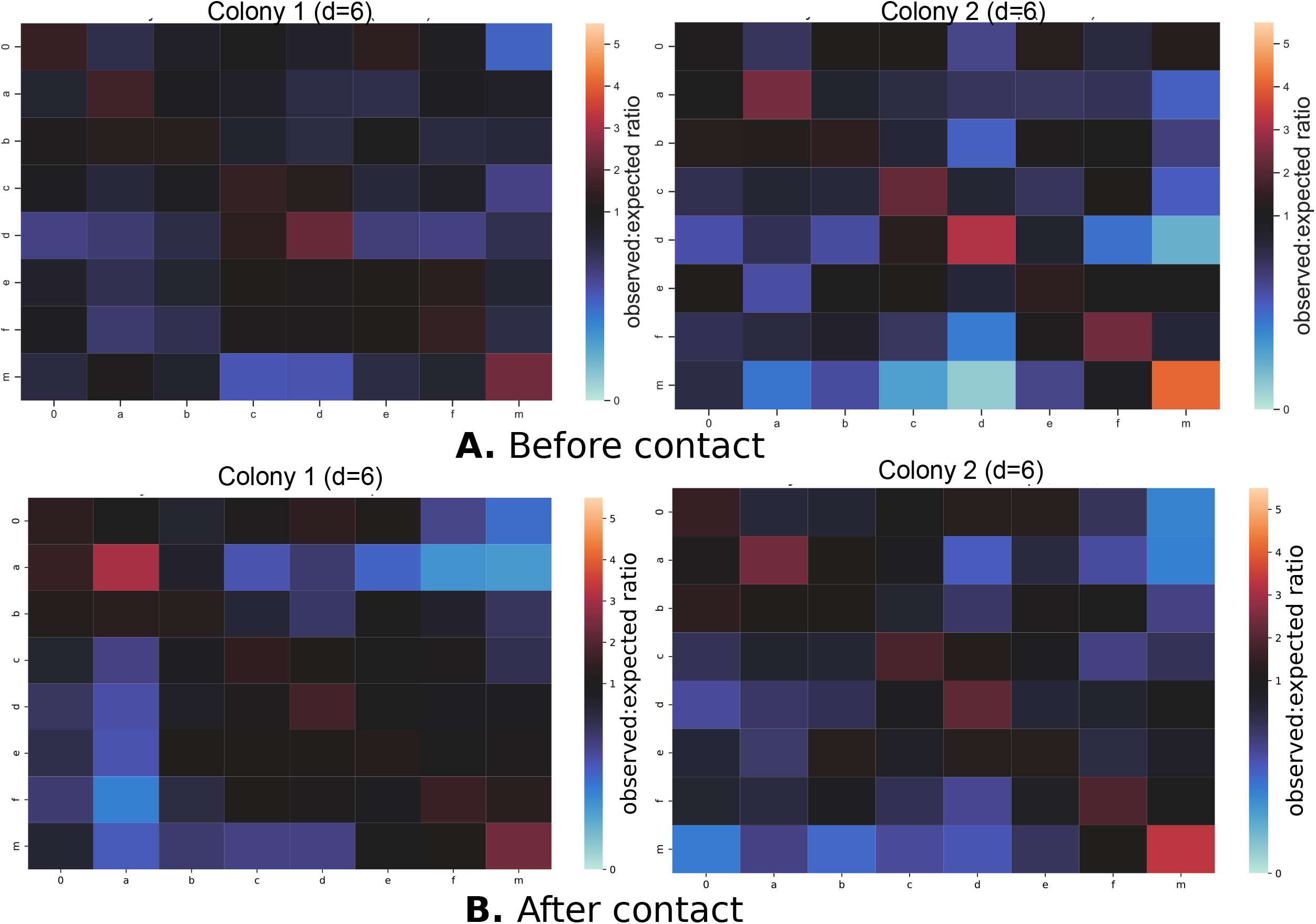
Ratio of observed to expected co-occurrence (d=6) of different note types in both the colonies before (**A**) and after (**B**) contact. Redder or warmer colors in the *ij-th* position in the matrix indicate that note type j occurred with note type i more often than expected by chance (see Methods for details). Note the higher propensity of note type d to co-occur with itself in Colony 2 (**A**), indicating an increased probability of repetition, which was lost after the colonies were brought into contact (**B**).

We examined this pattern further by quantifying how often different note types repeated themselves within warble songs across the two colonies. In the warbles of Colony 2, d notes exhibited an increased tendency to occur as repeats of 3 or more notes, and a decreased tendency to occur alone compared to Colony 1 (Fig 5A). After contact, however, males from the two colonies demonstrated broadly similar propensities to repeat all note types (Fig 5B). This pattern held true for each individual male in each colony; individual birds showed a pattern of note repetition specific to their colony of origin, which was lost after the colonies were brought into contact (Fig S5). Thus, our data were broadly consistent with repetitions of different note types (d notes in this particular case) acting as signatures of colony identity embedded in a complex and lengthy warble. After colonies came into contact, syntactic convergence caused these colony signatures to disappear, such that males from both erstwhile colonies produced warbles of similar structure, duration and complexity.

**Fig 5.**
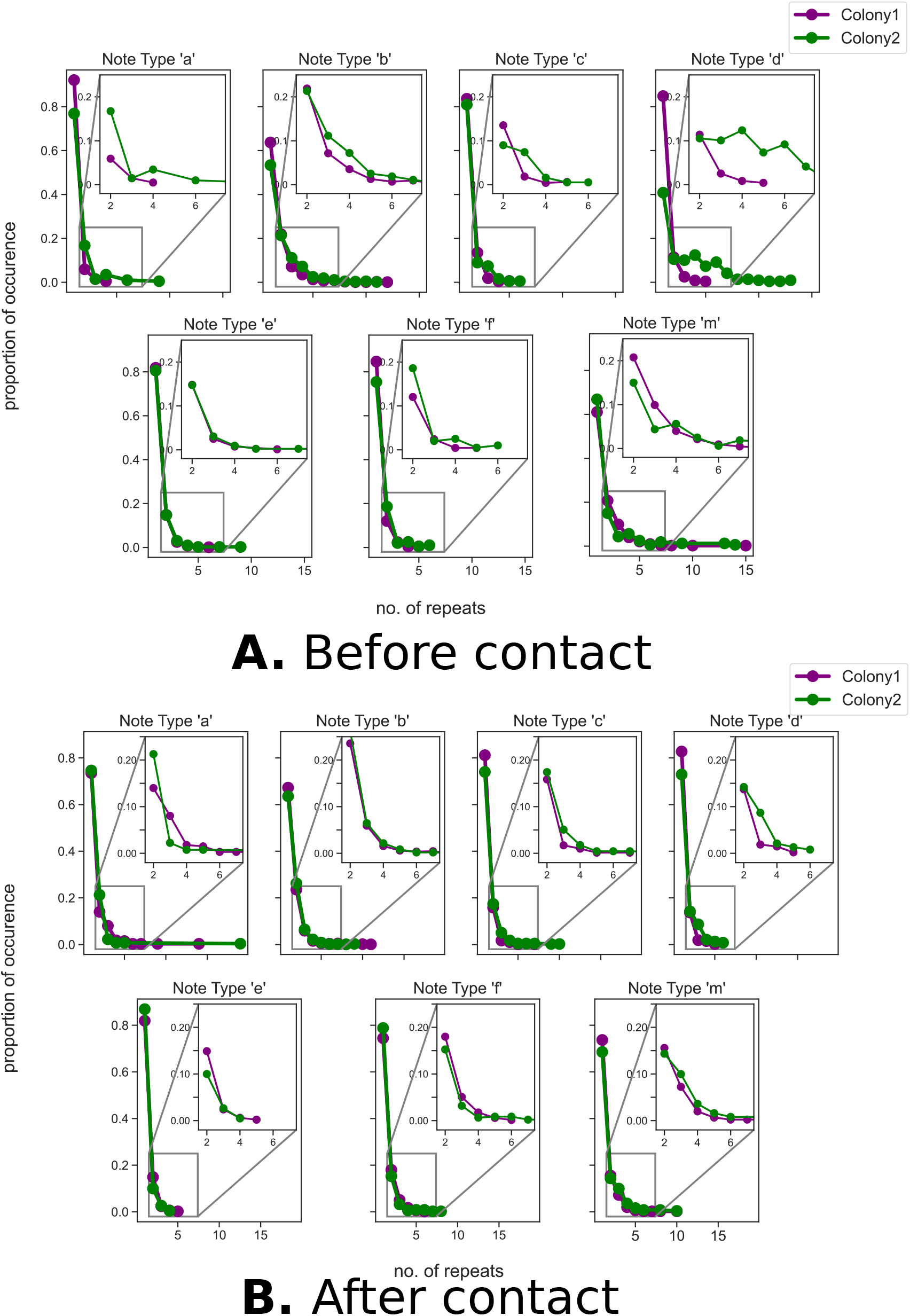
Proportion of occurrences of note repetitions of different lengths, for each note type for each colony before (**A**) and after (**B**) contact. Insets depict repeat lengths of 2 (e.g., a certain note type occurring twice consecutively) and greater, zoomed in for visual clarity. Males in Colony 2 were more likely to repeat d notes before contact, but this higher-order dialectic variation disappeared after the colonies were brought into contact.

## Discussion

Our study found that the warble song of budgerigars differed between individuals of different colonies (Fig 2, 3). Secondly, we found that these differences in complexity represent higher-order dialectic variation, resulting from an increased tendency of one colony to issue repetitive motifs of certain note types (Fig 4, 5). This, in turn, led to differing proportions of notes in vocal sequences across colonies (Fig 3), leading to differences in complexity as measured by the Shannon Entropy (Fig 2B). Our data thus suggest that the long, complex sequences of budgerigar warble songs contain embedded higher-order signatures of dialectic variation. When colonies come into contact, these differences are lost rapidly. A few months after contact, the syntactic structure of warble sequences had converged, and males from both erstwhile colonies sang warbles with similar syntactic structures,

Syntax in animal sensory signals is both diverse and complex, with considerable variety in its form and function (Arnold and Zuberbühler, 2006; Backhouse et al., 2022; Bhat et al., 2022; Briefer et al., 2013; Engesser et al., 2016; Engesser et al., 2019; Isaac and Marler, 1963; Kershenbaum and Garland, 2015; Kershenbaum et al., 2012; Kershenbaum et al., 2016; Leroux et al., 2021; Ligon et al., 2018; Scholes III, 2008). The structure of vocal sequences, particularly in vertebrates is known to convey information about context and behavioral state (Bhat et al., 2022; Ciaburri and Williams, 2019; Engesser et al., 2016; Leroux et al., 2021; Suzuki et al., 2018). For example, primates and pied babblers use different combinations of calls to convey different meanings (for example, different behavioral states) (Arnold and Zuberbühler, 2006; Clay and Zuberbühler, 2011; Engesser and Townsend, 2019; Engesser et al., 2016; Engesser et al., 2019) and chestnut-crowned babblers and hyraxes combine individually meaningless units into meaningful sequences that vary according to context (Engesser et al., 2019; Kershenbaum et al., 2012). On the contrary, we found that neither the duration nor complexity of budgerigar vocal sequences varied according to social context (Fig S2), that is, males produced similar warbles in both male-male and male-female contexts. Broadly, our results are concordant with another study suggesting that budgerigars sing similar warble songs to both male and female conspecifics (Tobin et al., 2019). Taken together, the available literature therefore suggests that warble song in budgerigars may serve a broader social function in communicating identity, similar to cetaceans and primates where acoustic signals serve to communicate group and individual identity (Candiotti et al., 2012; Deecke et al., 2010; Elowson and Snowdon, 1994; Mitani and Gros-Luis, 1998; Rendell and Whitehead, 2003).

The presence of unique, group-specific recognition signals may enable individuals to distinguish group members from non-group members (the password hypothesis) (Dahlin et al., 2014; Mammen and Nowicki, 1981; Tyack, 2008). Additionally, this similarity in call structure may also help mediate aggressive and affiliative interactions among group members (social association) (Dahlin et al., 2014; Mammen and Nowicki, 1981; Tyack, 2008). Most studies of parrots so far have suggested that these recognition signals are encoded in the acoustic structure of individual calls (Dahlin and Wright, 2012; Dahlin et al., 2014; Farabaugh et al., 1992; Hile and Striedter, 2000; Hile et al., 2000; Hile et al., 2005; Wanker and Fischer, 2001; Wanker et al., 2005). The structure of a flight call, for example, varies according to individual and colony (Dahlin et al., 2014; Farabaugh et al., 1992; Mammen and Nowicki, 1981). This acoustic structure (i.e. the time-frequency properties of an individual note) changes when individuals move between colonies or during pair-bonding and maintenance (Dahlin et al., 2014; Farabaugh et al., 1992; Hile and Striedter, 2000; Hile et al., 2000; Hile et al., 2005). A similar pattern (social vocal accommodation), has been described for the acoustic structure of social calls in primates (Elowson and Snowdon, 1994; Zürcher et al., 2021). However, individual notes represent only a single dimension of the complexity of parrot acoustic communication. Parrots, being perennial learners, frequently string together individual notes into complex vocal sequences that are also modified throughout their lifespans, often including mimicry of other sounds such as human speech (Balsby et al., 2017; Farabaugh et al., 1992; Pepperberg, 2009). Studies of intraspecific communication in parrots to date have focused almost exclusively on the acoustic properties of individual notes (Berg et al., 2012; Dahlin and Wright, 2009; Dahlin et al., 2014; Wright and Dahlin, 2017); very few studies have examined sequence structure or syntax in parrot vocalizations (Dahlin and Wright, 2009; Dahlin and Wright, 2012). We find that budgerigars from different social groups exhibit differences in organization of notes in a warble sequence, demonstrating higher-order dialectic variation. This variation is modified when groups come into contact, resulting in rapid syntactic convergence in just a few months (Fig 4, 5). Our data therefore suggest that budgerigars possess complex, multiscale encoding of group identity in their acoustic signals, and can modify this by social learning throughout their lifespan as flocks associate and dissociate in the wild. In nature, parrot social groups associate and dissociate into larger or smaller flocks in different seasons, and vocal convergence may help mediate social interactions as discussed earlier.

Our quantitative examination of signal structure and note co-occurrence patterns presents an example of the strength of computational approaches in studying animal communication (Bhat et al., 2022; Isaac and Marler, 1963; Kershenbaum and Garland, 2015). Evolutionary history and social structure of vocalizing vertebrate taxa influence both the acoustic properties of individual notes and their arrangement into syntactic structures (Arato and Fitch, 2021; Wanker et al., 2005; Wright and Dahlin, 2017). Vocal learning also exerts a strong influence on this process, and can lead to rapid divergence or convergence in acoustic signals (Fishbein et al., 2020; Garland et al., 2011; Lachlan and Servedio, 2004; Noad et al., 2000; Wilkins et al., 2013; Yeh and Servedio, 2015). By presenting evidence of higher-order dialectic variation and syntactic convergence in budgerigars, our study suggests a need to examine dialectic variation and communication of identity at multiple scales of signal organization. We also underscore the possibilities presented by psittaciform birds as model systems to examine the influence of complex social systems and social vocal learning on the structure and temporal organization of vocal sequences.

## Acknowledgments and Funding

We thank Ananda Shikhara Bhat for useful discussions and suggestions on analysis, Vaibhav Chhaya, Kunapareddy Kezia and Amey Danole for assistance during experiments, Prakash Raut and Suneel for bird care and Raghav Rajan and his lab for the use of their acoustic enclosures and for assistance and valuable discussions during the study. AK is funded by an INSPIRE Faculty Award from the Department of Science and Technology, Government of India, and an Early Career Research Grant (ECR/2017/001527) from the Science and Engineering Research Board (SERB), Government of India as well as an institutional initiation grant from IISER Bhopal. AJM and NW are recipients of the KVPY Fellowship, and PB of the INSPIRE Fellowship from the Government of India.

---

**Fig S1.** Examples of the different note types, in addition to those depicted in Fig 1.

**Fig S2.** Box plots depicting the duration and Shannon entropy of warbles across the two colonies before contact, sorted according to social context (male-male contexts versus male-female contexts). We did not observe significant differences in either warble duration or Shannon entropy across social contexts within a colony.

**Fig S3.** Ratio of observed to expected co-occurrence of different note types in both the colonies with d=3, before and after contact.

**Fig S4.** Ratio of observed to expected co-occurrence of different note types in both the colonies with d=9, before and after contact.

**Fig S5.** Proportion of occurrences of note repetitions of different lengths, for each note type, before and after contact. This is the same data as in Fig 5, but the lines represent each individual bird, color-coded according to colony identity.

