## Supplementary figures and images for "Higher-order dialectic variation and syntactic convergence in the complex warble song of budgerigars"

### Fig S1

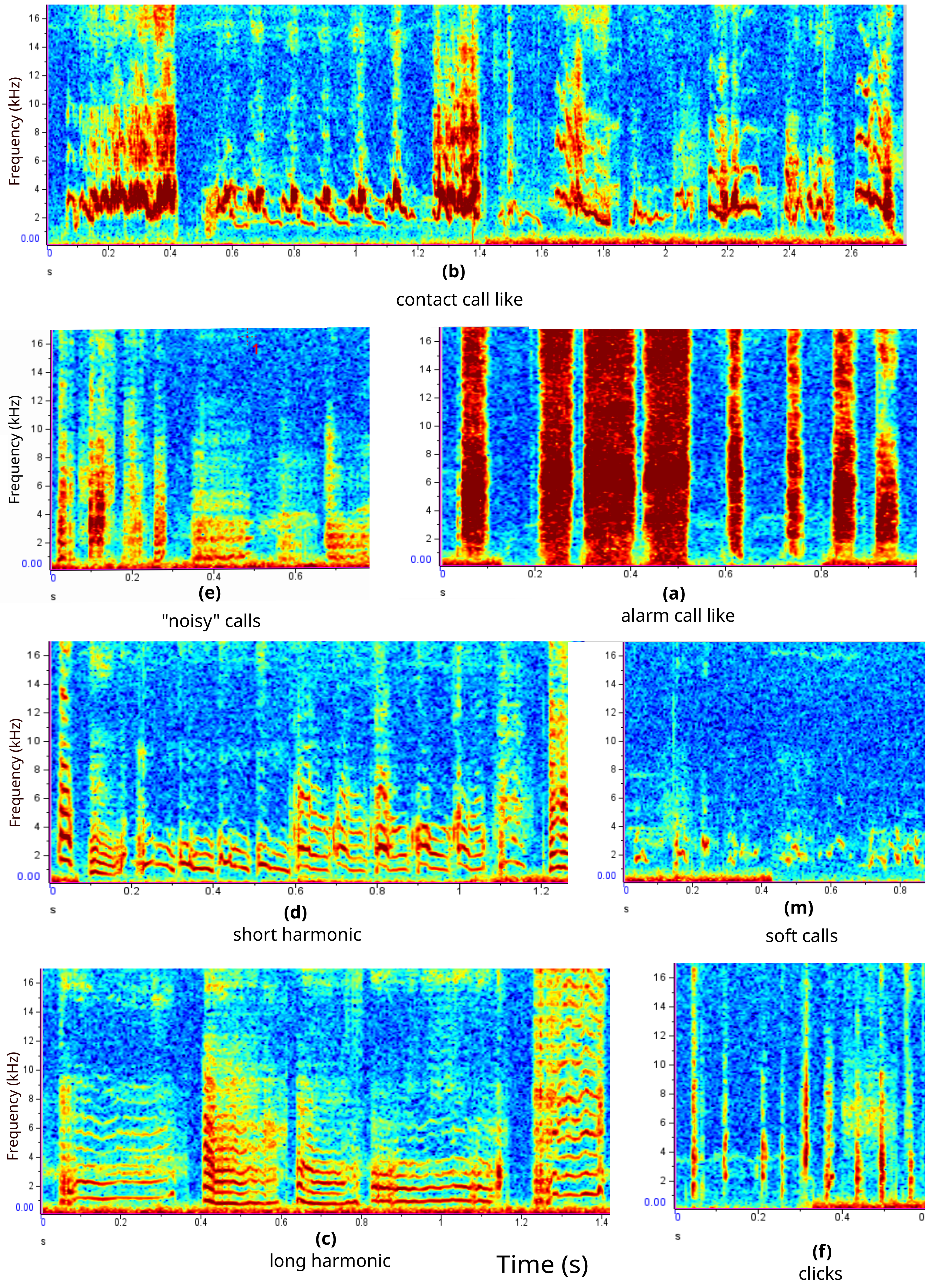

### Fig S2

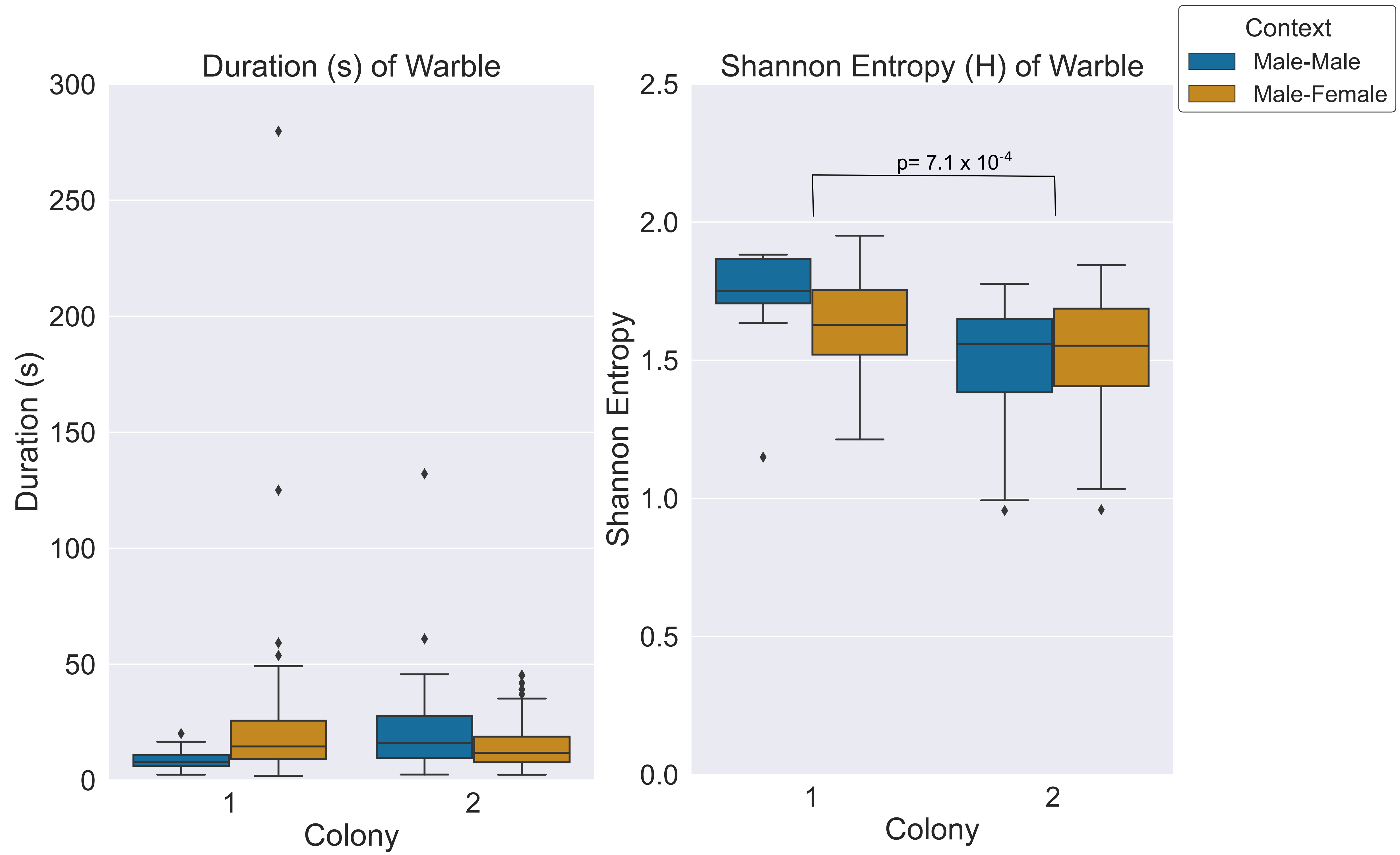

### Fig S3

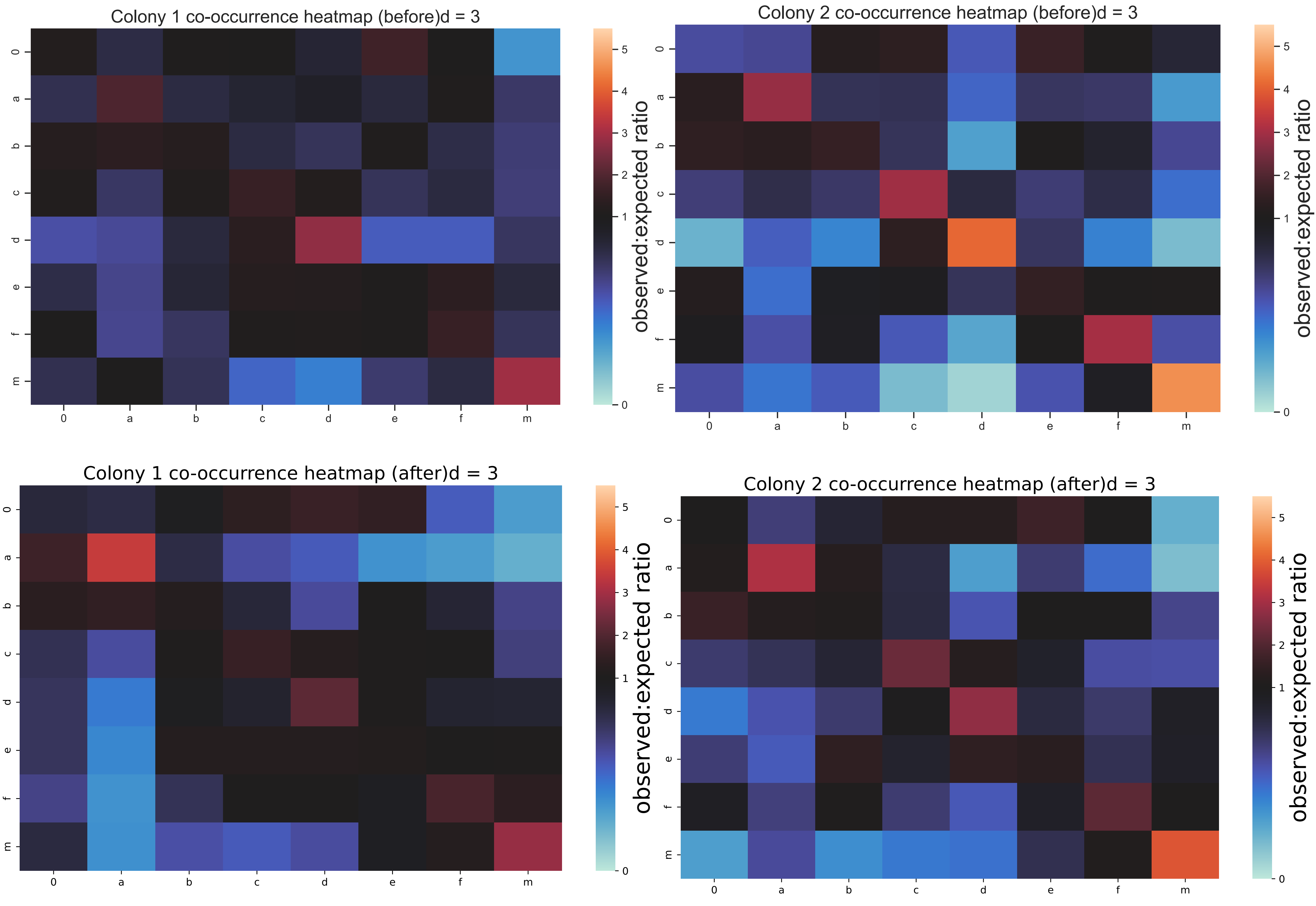

### Fig S4

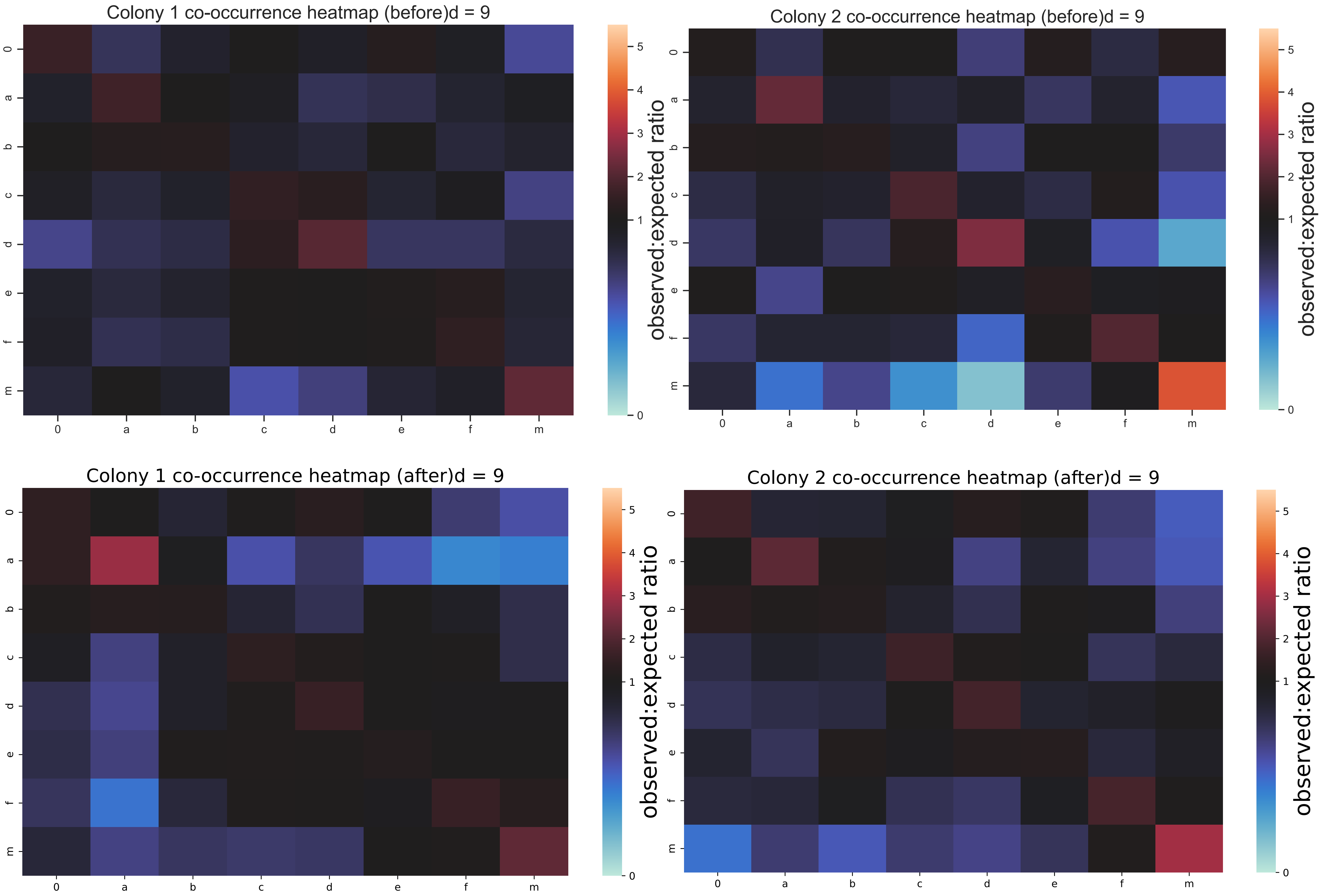

### Fig S5

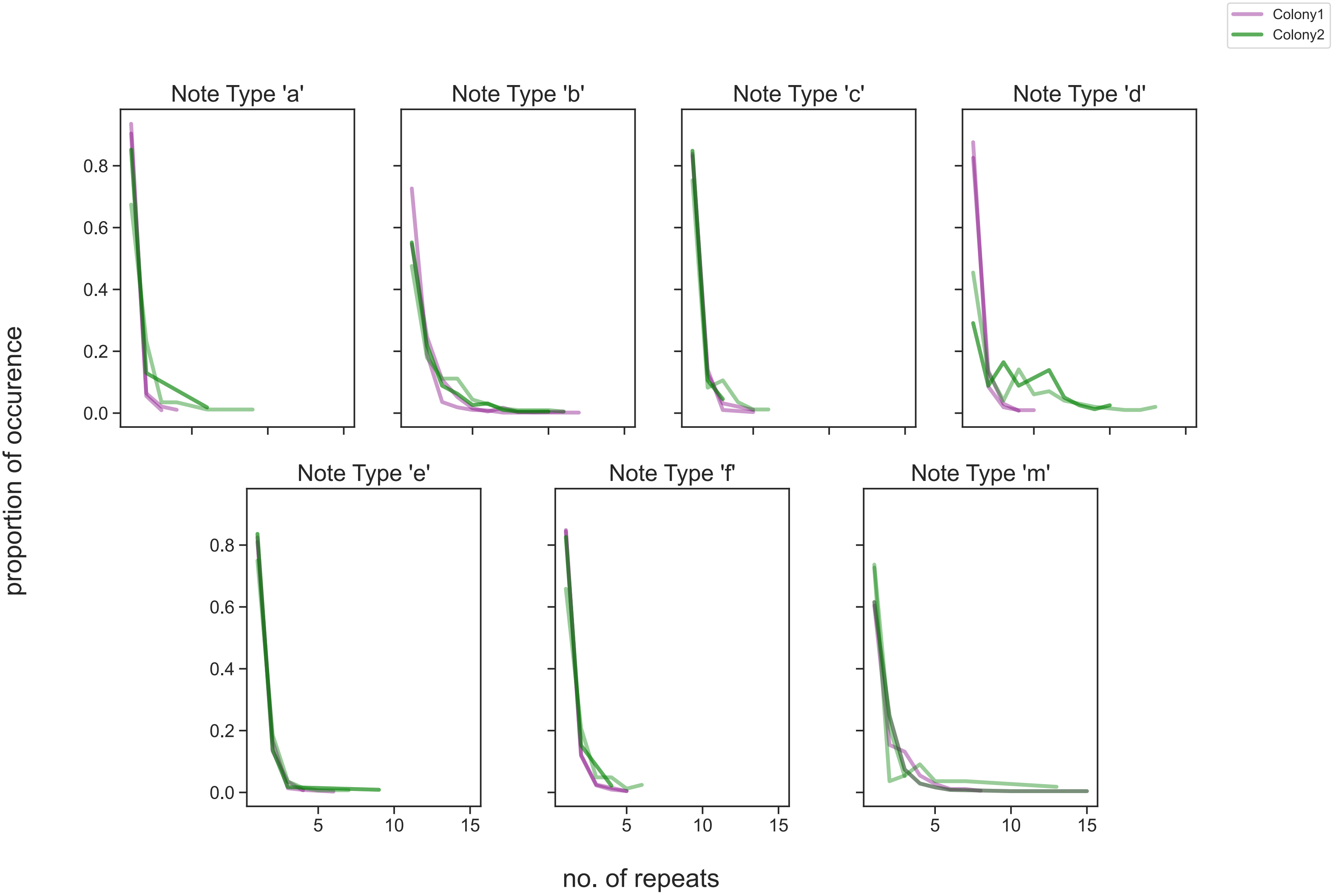


Before contact

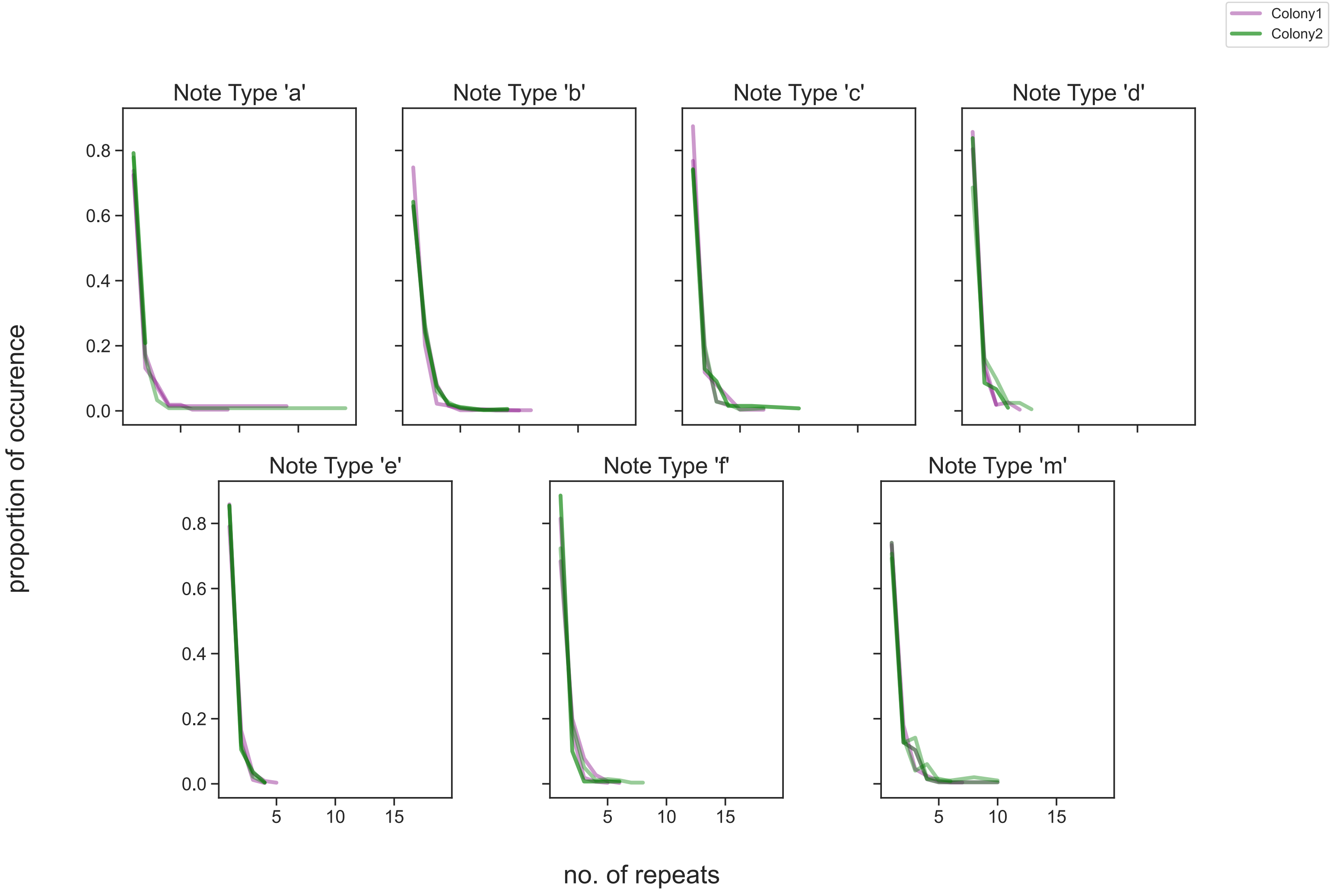


After contact
